# Swiprosin-1/EFhd2 promotes mitochondrial spare capacity in response to immobilized antigen in B cells via microtubule stabilization

**DOI:** 10.1101/2025.02.11.637645

**Authors:** Leonie Weckwerth, Philipp Tripal, Benjamin Schmid, Jana Thomas, Ann-Kathrin Himmelreich, Katharina Pracht, Sophia Urbanczyk, Hannah Allnach, Anna Gottwald, Ralph Palmisano, Dirk Mielenz

**Affiliations:** Division of Molecular Immunology, Department of Internal Medicine 3, Friedrich-Alexander-Universität Erlangen-Nürnberg and Universitätsklinikum Erlangen, Nikolaus-Fiebiger-Center, Glückstr. 6, 91054 Erlangen, Germany; Optical Imaging Center Erlangen (OICE), Friedrich-Alexander-Universität Erlangen-Nürnberg, Cauerstraße 3, 91058 Erlangen, Germany

**Keywords:** B cell, immune synapse, microtubules, mitochondria, spare capacity, EFhd2

## Abstract

B cells can recognize soluble and membrane bound antigens, enabling them to initiate and execute versatile immune responses. This study examines Swiprosin-1/EFhd2 (EFhd2) in regulating mitochondrial function and organization in B cells during B cell receptor (BCR) activation and immune synapse formation. Using EFhd2 knockout (KO) and wild-type (WT) murine B cells, we assessed mitochondrial abundance, membrane potential, and respiratory capacity using soluble anti-IgM and anti-CD40/IL-4 stimulation. EFhd2KO B cells exhibit more functional mitochondria and mitochondrial spare capacity selectively in activated but not in resting cells. This phenotype changed absolutely upon BCR synapse formation: While activated WT B cells enhance basal mitochondrial respiration, ATP production and maximal respiration, with large increments of spare capacity, EFhd2KO B cells fail completely to do so. Actin depolymerization and microtubule destabilization impair functional mitochondrial upregulation in WT but not in EFhd2KO B cells, while microtubule stabilization restores full spare capacity in EFhd2KO B cells at the BCR synapse. Live-cell imaging reveals that EFhd2KO B cells fail to organize mitochondria, microtubules, and BCRs symmetrically. Super-resolution 3D imaging shows that WT B cells condense mitochondria at the synapse, whereas EFhd2KO B cells display dispersed, unorganized mitochondria and BCR clusters. EFhd2 re-expression restores mitochondrial polarization in EFhd2KO B cells, confirming its role in coordinating mitochondrial positioning. These findings highlight EFhd2 as a key integrator of cytoskeletal and mitochondrial functions for optimal B cell responses to membrane bound antigens.

**Graphical Abstract:** 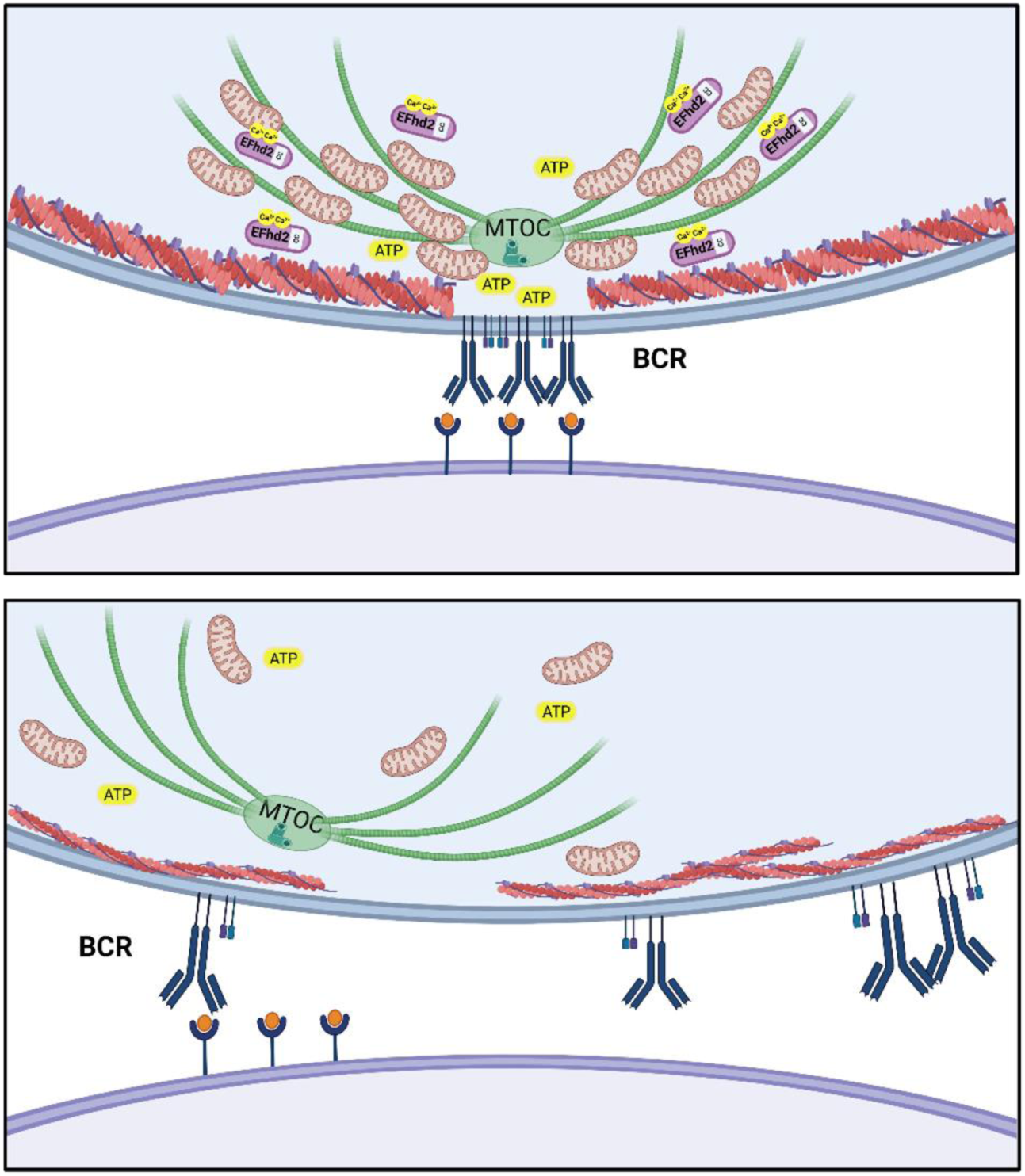

## Introduction

B cells can recognize a multitude of diverse soluble or membrane bound antigens owing to the unique structural organization of the B cell receptor (BCR) ^1^. Consequently, BCR activation can be mediated by antigen-presenting cells (APC), such as dendritic cells (DC), which present membrane bound antigens to B cells in the B cell follicle ^2,3^. Physiologically, repetitive immobile antigen arrays induce stronger anti-viral humoral immune responses than soluble antigen ^4,5^. After initial BCR activation, B cells migrate to the border of the B-T cell zone. Given cognate T cell help is available, B cells receive co-stimulatory signals through CD40 and the Interleukin-4 (IL-4) receptor, prompting them to migrate back to the B cell follicle where they can either seed germinal centers (GC) around the follicular DC (FDC) network ^6^, or undergo extrafollicular plasma cell differentiation. Over a period of 3-4 d, depending on the inciting antigen, early GC are being formed and mature GC are established after 7-10 d, ^7^. Mature GC contain two anatomically separate zones, the dark zone (DZ) and the light zone (LZ) ^8^. Repeated vaccination (booster) increases antibody (Ab) affinity over time ^9,10^ which occurs in GC via clonal expansion of B cells, followed by iterative cycles of somatic hypermutation (SHM) of Ab variable gene segments and B cell selection of GC B cells ^7^. To this end, FDC re-present antigen in the form of Ag-Ab immune complexes or complement-tagged antigens generated earlier. The LZ GC B cells recognize immobilized antigen on FDC via their BCR and specialized synapses owing to LZ B cell and FDC intrinsic features ^11,12,13^. These synapses function to assess the BCRs for preserved and, at best, improved functionality ^14–16^. B cells test the affinity of their mutated BCR to antigen immobilized on FDC by exerting force via coupling of the BCR to the Actin-myosin system ^11^. Force generation in the (F)DC-B cell synapse is proportional to BCR affinity and will favor high affine B cells to acquire, internalize and present antigen to follicular T helper cells (TFH) ^15^. Thereby, the LZ B cells receive positive selection signals via BCR signaling^17^ and co-stimulation that can elicit MYC expression and anabolic pathways, such as the mammalian target of Rapamycin (mTOR) pathway ^18^. This signaling also promotes recycling to the DZ ^18–20,21^ or, finally, GC exit, plasma cell and memory B cell (MBC) maturation. Notably, catabolic pathways control GC B cell biology as well. Thus, in particular pre-GC B cells as well as the early GC phase demand glycolysis and lactate production by lactate dehydrogenase ^22–24^. Conversely, in vitro activated GC B cells from BCR transgenic mice transferred into antigen-unresponsive mice prefer fatty acid oxidation in mitochondria ^25^. In alignment, regulation of genes required for oxidative phosphorylation (OxPhos) in LZ GC B cells, as well as transcription factor A (TFAM) activity, support GC responses and antibody affinity maturation dynamically ^26–28^. Of note, catabolic pathways can be an important upstream prerequisite of anabolic mTOR activity in activated B cells ^22,28^. Interestingly, mitochondrial mass and their spatial organization are more pronounced in LZ than in DZ GC B cells, suggesting that the location of GC B cells and their contact with FDC control respiratory chain activity ^26^. This would be in line with BCR affinity-coupled OxPhos gene expression ^27^. Together, these data suggest that the organization of the LZ B-FDC synapse via the BCR directs mitochondrial function.

Previously we have shown that the speed and velocity of GC B cells are proportional to GC plasma cell output ^29^. To show this, we have used B cells from mice deficient for the Actin-binding and bundling protein Swiprosin-1/EFhd2 (EFhd2) ^30,31^. EFhd2’s Actin-bundling activity is enhanced by binding of Ca^2+^ to the EF-hands ^32,33^. Besides binding F-Actin, EFhd2 also interacts with the microtubule (MT) system by slowing down kinesin-mediated anterograde transport on microtubules ^34^. Loss of EFhd2 increases the speed of BCR-activated B cells in GC in vivo as well as on intercellular adhesion molecule (ICAM) 1 and CXCL13 coated slides, by increasing Actin dynamics at the expense of B cell polarization ^30^. Hence, the GC plasma cell output and IgG and IgE antibody level is higher in EFhd2KO mice ^29,30^. Unexpectedly, IgG antibody affinity towards the hapten Nitrophenol (NP), as determined by ELISA, is not elevated concomitantly, suggesting a LZ phenotype. In fact, EFhd2KO B cells form smaller and less organized synapses with FDC in vivo, as shown by 2-Photon microscopy ^29^, and they have competitive disadvantages in the GC^29^. Overexpression of EFhd2 supports the inherent apoptosis induced by soluble anti-IgM antibodies in WEHI231 as well as loss of the mitochondrial membrane potential (Ψ), while knock-down of EFhd2 augments survival under these conditions ^35^. These data point to a connection of EFhd2 to the BCR-mitochondrial axis which controls B cell survival ^36^. Thus, we hypothesized that EFhd2 modulates mitochondrial function in response to soluble or membrane-bound BCR engagement. Here, we show that primary activated EFhd2^+/+^ (wildtype; WT) cells strongly increase mitochondrial function when attached to immobilized anti-IgM antibodies. EFhd2KO B cells fail completely to do so due to defective BCR and mitochondrial organization at the B cell synapse, but microtubule stabilization restores mitochondrial function. Hence, the BCR can sense antigenic nature via EFhd2 mediated microtubule stabilization and translate it into mitochondrial fitness.

## Results

### EFhd2 expression in GC B cells impacts on functional mitochondria

The proportional relation between BCR-elicited mitochondrial apoptosis and EFhd2 abundance in WEHI231 cells^35^ suggested that EFhd2 controls mitochondrial viability in response to BCR activation elicited by soluble antigen. On the other hand, EFhd2 might link mitochondrial function to BCR synapse formation. Assuming that some isolated GC B cells had recent contact with FDC, we analyzed mitochondrial abundance and ψ in GC B cells of mesenteric lymph nodes (mLN) of WT and EFhd2^−/−^ (EFhd2 knock-out, KO) mice by flow cytometry (Figure 1A, B; gating strategy in Figure S1). Mitochondria were classified as functional (Mitotracker Deep Red^high^, Mitotracker Green^high^ or ^low^) or dysfunctional (Mitotracker Deep Red^low^, Mitotracker Green^pos^) (Figure 1B). While there was no sign of mitochondrial alteration in CD19^+^CD38^−^GL7^+/low^CD95^+^ GC B cells (Figure 1B, C, D), CD19^+^CD38^−^GL7^−^CD95^+^ EFhd2KO cells, defined as late GC B cells ^37^, exhibited a reduced ψ (Figure 1E); however, resting non GC B cells showed no alterations (Figure 1F).

**Figure 1:**
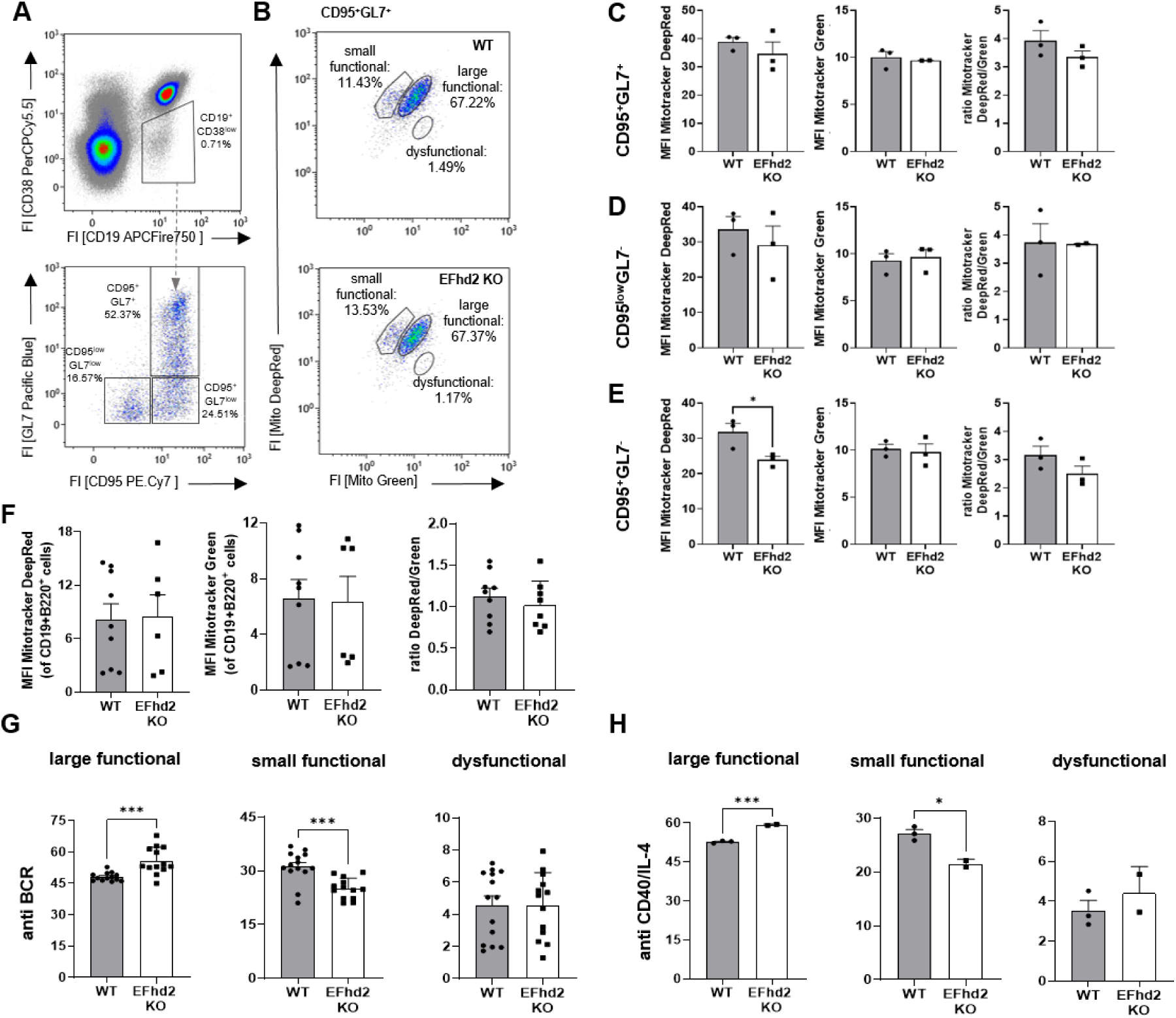
Effect of EFhd2 on mitochondria in B cells. A. Mesenteric lymph node cells (mLN) from WT and EFhd2KO mice were stained as indicated, as well as with Mitotracker Green (Mito Green) and Mitotracker Red (Mito Red), and analyzed by flow cytometry. Germinal center (GC) B cells were defined as CD19^+^CD38^low^ and further via anti CD95 and GL7 staining. Representative plots are depicted. Numbers indicate frequencies in the indicated gates. B. Analysis of mitochondrial size and membrane potential of CD95^+^GL7^+^ GC B cells. C-E. Mean fluorescence intensity (MFI) of Mito Green and Mito Red of the indicated GC populations. F. MFI of Mito Green and Mito Red of CD19^+^B220^+^CD38^+^ non-GC populations. G-H. splenic B cells from WT and EFhd2 KO mice were activated with soluble anti-IgM Ab or anti-CD40 Ab + IL-4 for 24h. Frequencies of large functional (Mitotracker Deep Red^high^, Mitotracker Green^high^), small functional (Mitotracker Deep Red^high^, Mitotracker Green^low^) and dysfunctional (Mitotracker Deep Red^low^, Mitotracker Green^pos^) of splenic B cells after 24h of anti-IgM (G) or anti-CD40 + IL-4 (H) activation are depicted. Data are presented as mean ± SEM; each dot represents one mouse; statistics; pooled from 1-5 experiments. Significance was analyzed using Shapiro-Wilk and unpaired t-test (p values: *p < 0.05, **p < 0.01, ***p < 0.002).

GC B cells receive alternating or simultaneous signals via tonic BCR signaling, membrane bound antigen, CD40, the IL-4 receptor and others ^38^. To segregate the multiple signals, we activated primary murine splenic WT and EFhd2KO B cells via sole anti-IgM stimulation for 24h. The Mitotracker Deep Red^high^, Mitotracker Green^high^ population was larger than the Mitotracker Deep Red^high^, Mitotracker Green^low^ population in activated EFhd2KO B cells (Figure 1G), whereas the proportion of cells with dysfunctional mitochondria was unchanged in primary EFhd2KO B cells. Similar results were obtained upon anti CD40/IL-4 stimulation (Figure 1H). Thus, EFhd2 impacts mitochondrial mass rather than apoptosis in primary B cells, and seems to have opposing functions in late GC than in primary B cells activated by soluble stimuli.

### EFhd2 switches mitochondrial spare capacity in soluble versus plate-bound anti-IgM stimulated B cells

To see whether EFhd2 would impact mitochondrial function under immune-synapse simulating conditions, we continued to work with shortly activated B cells because a) non-activated B cells are metabolically inert ^39^, b) sorted GC B cells would be too short in number and viability and c) are heterogeneously imprinted by various signals. Hence, we analyzed 24h soluble anti-IgM activated cells attached to Poly-D-Lysine (PDL) by extracellular flux analysis via a Seahorse Mito Stress Test (MST) (Figure 2). In a MST, cells respire basally. Oligomycin stops ATP Synthase and O_2_ consumption, which will in turn increase to its maximum via uncoupling reagents such as Carbonyl cyanide-p-trifluoromethoxyphenylhydrazone (FCCP). O_2_ consumption will cease again after the abolishment of complex I and III activity by adding Rotenone/Antimycin A (Figure 2A). The MST performed with 24h soluble anti-IgM activated cells showed that EFhd2KO B cells displayed a tendency towards higher mitochondrial activity (Figure 2B, D, E), which correlates with the higher proportion of cells containing more mitochondria (Figure 1G). To simulate an immune synapse, we subsequently coated the Seahorse assay plates next with anti-IgM antibodies. Strikingly, all mitochondrial parameters of WT B cells increased on anti-IgM antibodies, while EFhd2KO B cells did not react at all (Figure 2C-E). This difference was not due to limiting coating concentrations (Figure S2A, B) or altered viability (Figure S2C). Also after soluble anti-CD40/IL-4 stimulation, only WT B cells strongly upregulate mitochondrial spare capacity (SPRC) while EFhd2KO B cells are completely refractory, despite trends towards higher basal SPRC (Figure 2H), basal respiration, ATP production and proton leak (Figure S2C). This could be due to increased mitochondrial mass, but not increased mTOR activity (Figure S3A-D). Together, these experiments reveal that i) activated B cells increase mitochondrial function upon membrane bound antigen contact and ii) require EFhd2 for this process.

**Figure 2:**
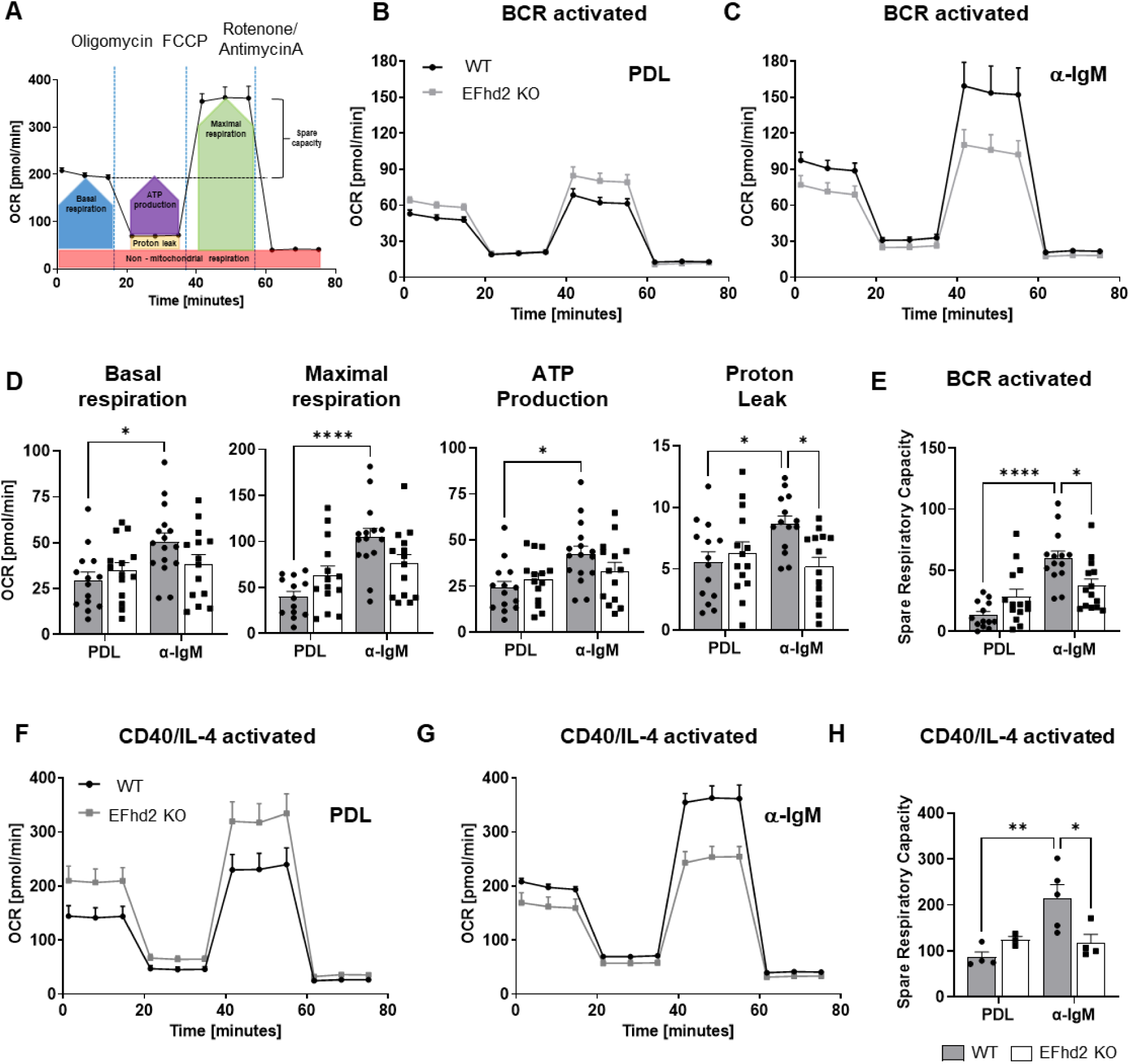
EFhd2 increases mitochondrial function upon immobilized BCR activation. A. Schematic of the experimental setup. B. Representative extracellular flux analysis of 24h anti-IgM Ab activated splenic B cells attached to poly-D-lysine (PDL) or anti IgM antibodies (C). The basal oxygen consumption rate (OCR) was measured before and after injection of oligomycin, FCCP and Rotenone plus Antimycin A using the Seahorse Wave Mito Stress Test (MST) protocol. Symbols represent means of two mice, each of three to four replicate wells, mean ± SEM. D. Calculated OCR of basal respiration, maximal respiration, ATP production and proton leak of 24h anti-IgM activated B cells attached to PDL or anti IgM antibodies. E. Calculated spare respiratory capacity (SPRC). D, E: data are presented as mean ± SEM; each dot represents one mouse; statistics: pooled from 7-8 experiments with 2-3 mice per genotype per experiment; significance was analyzed using Two-way ANOVA (p values <0.5. *p < 0.05, **p < 0.01, ***p < 0.002, ****p < 0.0004). F, G. Representative extracellular flux analysis of 24h anti-CD40/IL-4 activated splenic B cells attached to PDL or anti IgM antibodies. The basal oxygen consumption rate (OCR) was measured before and after injection of oligomycin, FCCP and Rotenone plus Antimycin A using the Seahorse Wave Mito Stress Test (MST) protocol. Symbols represent means of two mice, each of three to four replicate wells, mean ± SEM. H. Calculated SPRC. Data are presented as mean ± SEM; each dot represents one mouse; pooled from 1-2 experiments with each 3-5 independent cultures per genotype; significance was analyzed using Two-way ANOVA (p values <0.5. *p < 0.05, **p < 0.01, ***p < 0.002, ****p < 0.0004).

### B cells increase mitochondrial spare capacity on immobilized anti-IgM antibodies via the Actin cytoskeleton

EFhd2 stabilizes Actin filaments and activated EFhd2KO B cells exhibit a more dynamic Actin cytoskeleton ^29^. Immune synapse formation depends on Actin remodeling ^12^. To test whether the Actin cytoskeleton is involved in connecting the BCR to mitochondria on immobilized antigen we treated WT and EFhd2KO B cells with the Actin-depolymerizing drug Latrunculin A (LA). While LA reduced maximal respiration significantly in WT B cells, there was no significant effect in EFhd2KO B cells (Figure 3A). Although WT B cells were treated with LA, they still increased basal respiration, ATP production and proton leak (Figure 3B, C). Nevertheless, LA reduced SPRC in BCR attached WT B cells while aligning SPRC on PDL to levels comparable to those observed in EFhd2KO B cells (Figure 3D). This experiment shows that proper Actin dynamics contributes to the BCR-mitochondrial axis. It also confirms that the Actin cytoskeleton is already less stable in EFhd2KO B cells because LA is ineffective and suggests that Actin depolymerization could boost mitochondrial activity.

**Figure 3:**
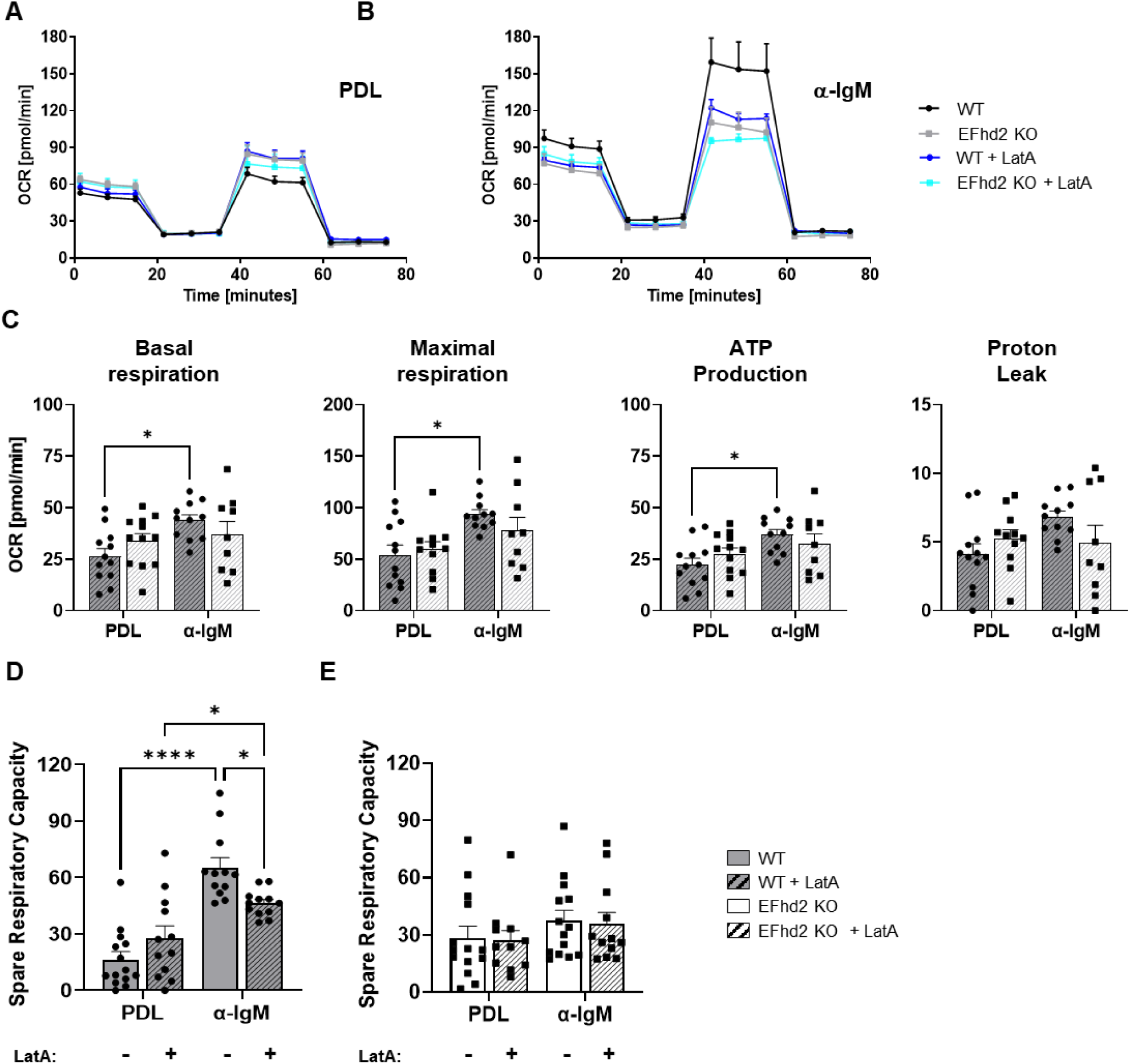
Latrunculin A decreases mitochondrial function upon immobilized BCR activation. A, B. Representative extracellular flux analysis of 24h anti-IgM Ab activated splenic B cells attached to poly-D-lysine (PDL) or anti IgM antibodies with or without Latrunculin A (Lat A) treatment. The basal oxygen consumption rate (OCR) was measured before and after injection of oligomycin, FCCP and Rotenone plus Antimycin A using the Seahorse Wave Mito Stress Test (MST) protocol. Symbols represent means of two mice, each of three to four replicate wells, mean ± SEM. C. Calculated OCR of basal respiration, maximal respiration, ATP production and proton leak of 24h anti-IgM activated B cells attached to PDL or anti IgM antibodies antibodies with or without Latrunculin A (Lat A) treatment. D, E. Calculated SPRC. C-E: Data are presented as mean ± SEM; each dot represents one mouse; pooled from 6-8 experiments with each 1-3 independent cultures; significance was analyzed using Two-way ANOVA (p values <0.5. *p < 0.05, **p < 0.01, ***p < 0.002, ****p < 0.0004).

### B cells increase mitochondrial spare capacity on immobilized anti-IgM antibodies via the tubulin cytoskeleton

Besides the Actin system, microtubules are also involved in immune synapse organization but not many data in B cells are available ^40^. EFhd2 interferes with Kinesin-mediated transport along microtubules in vitro ^34^. Moreover, EFhd2KO B cells appear to loose polarity during cell migration, a process that is coordinated by microtubule dependent anterograde transport ^40,41^. Therefore, activated EFhd2WT and KO B cells were attached to anti-IgM antibodies in the presence of Nocodazole, which depolymerizes microtubules ^42^. In comparison to LA, Nocodazole hindered the prominent increase of basal and maximal respiration, ATP production, proton leak and SPRC (Figure 4A-D) while elevating SPRC in WT B cells attached to PDL. Strikingly, Nocodazole did not exert any effect on EFhd2KO B cells. This demonstrates that both EFhd2 as well as stabilization of the microtubule cytoskeleton are essential for the BCR-mitochondrial axis. To challenge this notion, we chose an opposite approach and applied Paclitaxel (PTX) to stabilize microtubules ^43^. Interestingly, also PTX partially prevented the increase of basal and maximal respiration, ATP production and proton leak in BCR attached WT B cells, while there was a less pronounced effect on EFhd2KO B cells (Figure 5A-C). However, Paclitaxel did not modulate SPRC in BCR attached WT B cells (Figure 5D). In stark contrast, Paclitaxel fully restored SPRC of EFhd2KO B cells upon attaching to anti-IgM antibodies (Figure 5E). Noteworthy, none of the used Actin or tubulin-modulating pharmaceuticals affected viability (Figure S3E-G). Together, these data unequivocally demonstrate that heightened SPRC elicited by plate-bound BCR engagement depends on EFhd2-mediated microtubule stabilization.

**Figure 4:**
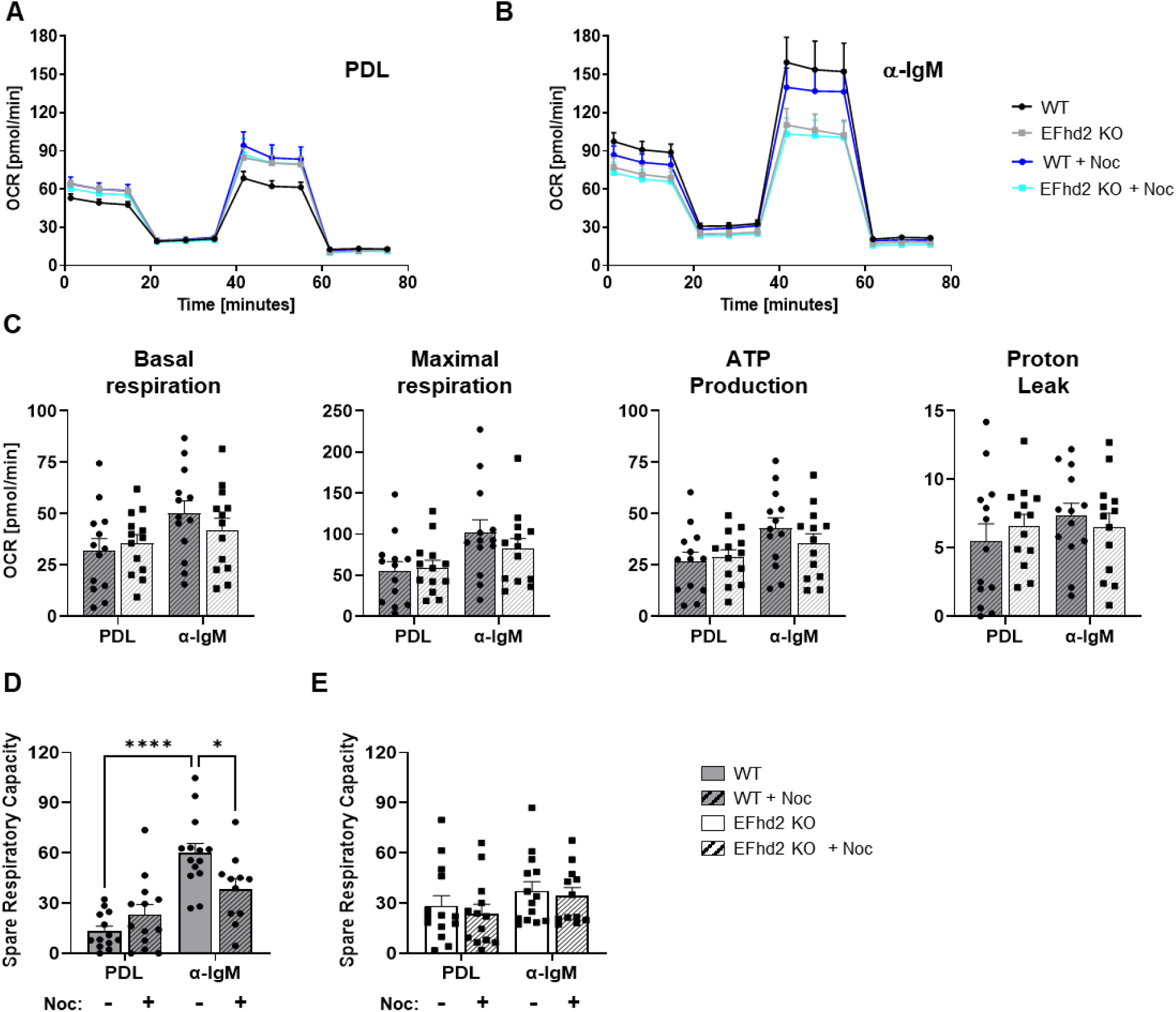
Microtubule destabilization decreases mitochondrial function upon immobilized BCR activation. A, B. Representative extracellular flux analysis of 24h anti-IgM Ab activated splenic B cells attached to poly-D-lysine (PDL) or anti IgM antibodies with or without Nocodazole (Noc) treatment. The basal oxygen consumption rate (OCR) was measured before and after injection of oligomycin, FCCP and Rotenone plus Antimycin A using the Seahorse Wave Mito Stress Test (MST) protocol. Symbols represent means of two mice, each of three to four replicate wells, mean ± SEM. C. Calculated OCR of basal respiration, maximal respiration, ATP production and proton leak of 24h anti-IgM activated B cells attached to PDL or anti IgM antibodies antibodies with or without Nocodazole treatment. D, E: Calculated SPRC. Data are presented as mean ± SEM; each dot represents one mouse; pooled from 6-8 experiments with each 1-3 independent cultures; significance was analyzed using Two-way ANOVA (p values <0.5. *p < 0.05, **p < 0.01, ***p < 0.002, ****p < 0.0004).

**Figure 5:**
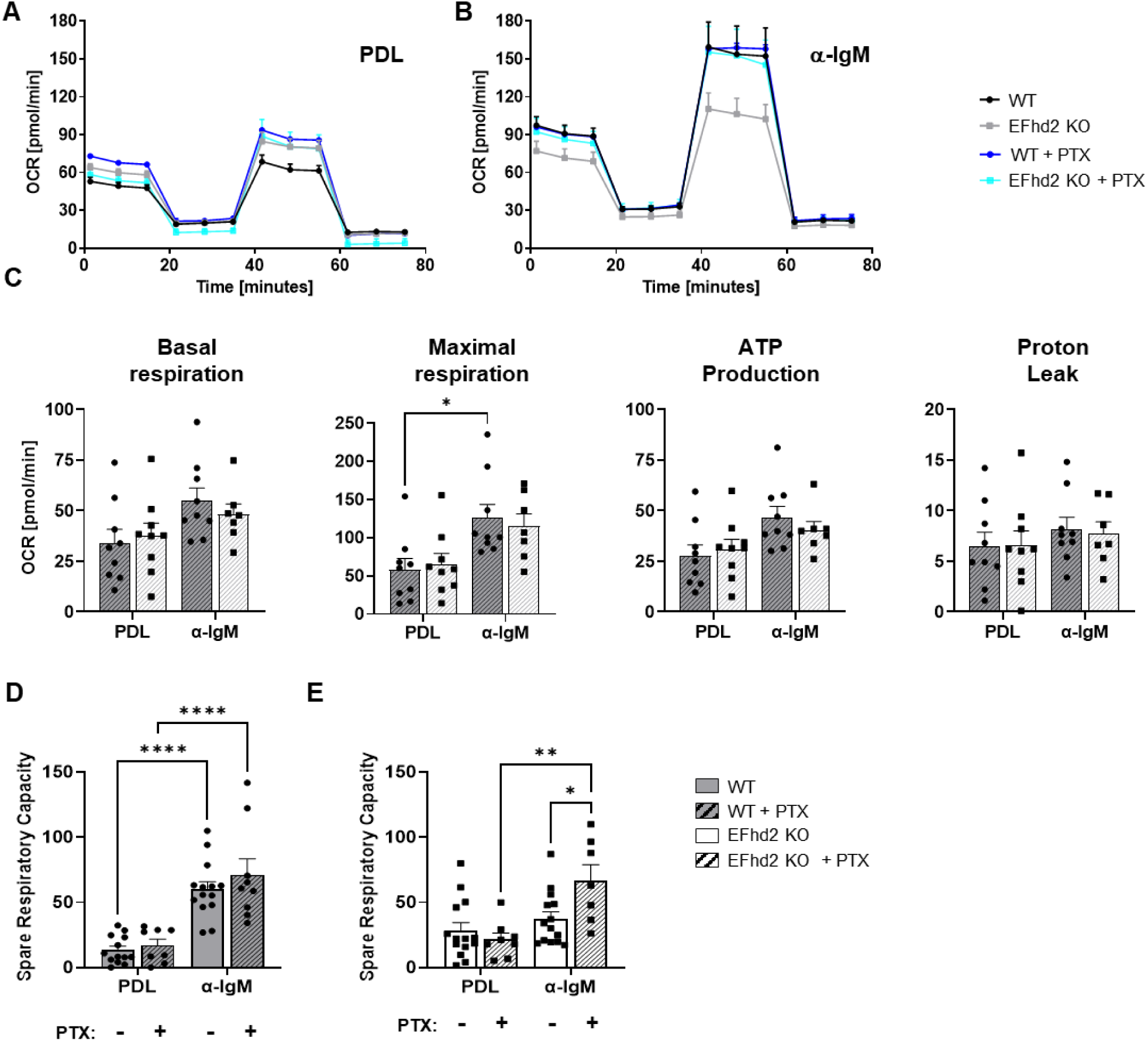
EFhd2-dependent microtubule stabilization drives mitochondrial function upon immobilized BCR activation. A, B. Representative extracellular flux analysis of 24h anti-IgM Ab activated splenic B cells attached to poly-D-lysine (PDL) or anti IgM antibodies with or without Paclitaxel (PTX) treatment. The basal oxygen consumption rate (OCR) was measured before and after injection of oligomycin, FCCP and Rotenone plus Antimycin A using the Seahorse Wave Mito Stress Test (MST) protocol. Each Symbols represent means of two mice, each of three to four replicate wells, mean ± SEM. C. Calculated OCR of basal respiration, maximal respiration, ATP production and proton leak of 24h anti-IgM activated B cells attached to PDL or anti IgM antibodies antibodies with or without Paclitaxel treatment. D, E. Calculated SPRC. Data are presented as mean ± SEM; each dot represents one mouse; statistics: pooled from 4-8 experiments with each 1-3 independent cultures; significance was analyzed using Two-way ANOVA (p values <0.5. *p < 0.05, **p < 0.01, ***p < 0.002, ****p < 0.0004).

### EFhd2KO B cells fail to organize mitochondria, microtubules and the BCR symmetrically in the synapse

Owing to the microtubule-associated control of SPRC in our system, a correlation of EFhd2 with mitochondrial morphology seemed likely. Consequently, we visualized mitochondria, microtubules and the BCR in live cells with Mitotracker Green, Tubulin tracker Deep Red and Alexa 647 labeled anti IgM Fab fragments. The activated B cells were adhered to PDL or anti-IgM coated glass slides and analyzed by spinning disc live cell microscopy. Representative 3D reconstructions reveal that WT B cells symmetrically organize BCR clusters in vicinity to an Aster-like microtubule and mitochondria network (Figure 6A). This tethers the mitochondria thereby to the B cell-substrate interface. In contrast, the microtubule network in EFhd2KO B cells is displaced more asymmetrically; the mitochondrial network is less well aligned the microtubule organization; and BCR clusters are more dispersed and disconnected. To quantify these findings, we defined the microtubule organizing center (MTOC) as an aggregation of tubulin at the cell-substrate interface from which the Aster-like microtubule network spreads out (Figure 6A). We then measured the distance of the MTOC to the focus of the cells on PDL and on anti-IgM antibodies. While there was no difference in cellular polarization in cells imaged on PDL (Figure 6B), the distance of the MTOC to the focus of the cells was higher in BCR-bound EFhd2KO than in WT B cells (Figure 6D). These data demonstrate that in response to attachment of B cells via the BCR, EFhd2 serves as a factor for proper B cell polarization. If this was due to a distorted microtubule-BCR connection, the MTOC would be expected to distribute randomly over the cells. However, we did observe a polarization to the basal side of the cells. Therefore, we calculated the weighted distance of the BCR to the MTOC, which was larger in Ehd2KO B cells regardless of the substrate (Figure 6B, C). As the BCR appears to have lost track without EFhd2, we suspected a smaller mitochondrial area in vicinity to the cell-substrate contact area. In fact, the focal mitochondrial area was smaller in BCR-attached EFhd2KO B cells on anti-IgM antibodies, but not on PDL (Figure 6D).

**Figure 6:**
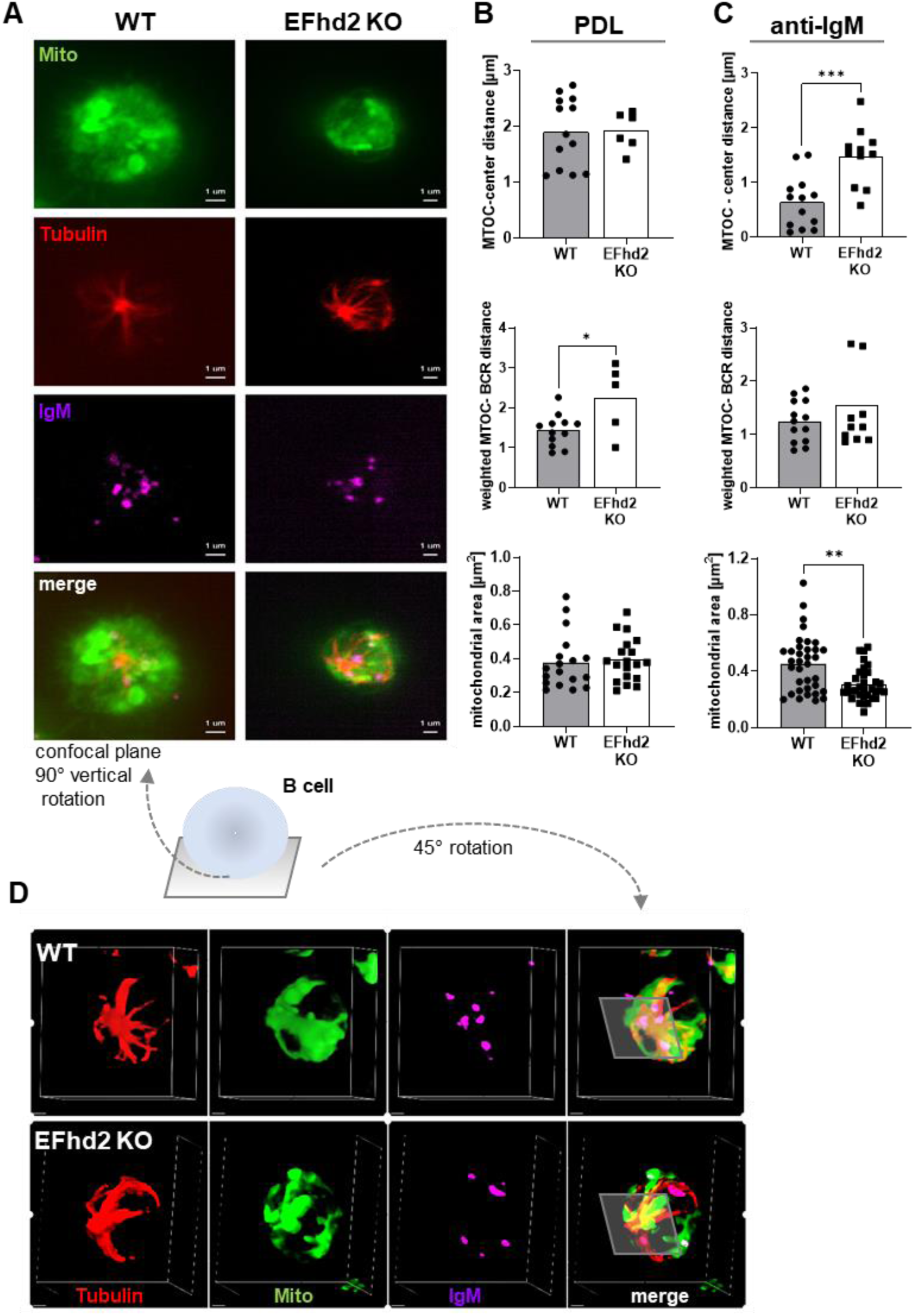
EFhd2 controls mitochondrial abundance at the BCR synapse. A. representative spinning disc live cell images of one 24h anti IgM Ab activated WT (left) or EFhd2 KO (right) B cells attached to anti IgM Ab. Mitochondria, microtubules and the BCR were visualized in live cells with Mitotracker Green, Tubulintracker DeepRed and Alexa 647 labeled anti-IgM Fab fragments, respectively. Fluorescence of the mitochondria (green), tubulin (red) and the BCR (magenta) are shown both separately and merged (scale bar, 1µM). B, C. Cell polarization (top), weighted distance between the microtubule organizing center (MTOC) and the BCR (middle), or mitochondrial area (bottom), were assessed directly after attachment on poly-D-Lysine (PDL) or anti-IgM Ab. D. 45° rotated 3D view from WT (upper panel) and EFhd2 KO (lower panel) shown in a. data are presented as mean ± SEM; each dot represents one cell. Significance was analyzed using Shapiro-Wilk and unpaired t-test (p values: *p < 0.05, **p < 0.01, ***p < 0.002).

### EFhd2 controls BCR and mitochondrial reorganization at the BCR synapse

To correlate the morphology to the EFhd2-controlled SPRC, we reconstructed the mitochondrial network by super resolution imaging to enable the view of 3D reconstructed cells from different angles (Figure 7). CD40/IL-4 activated WT B cells attached to PDL exhibited structured mitochondrial and BCR networks which, however, did not emerge to be well connected. The mitochondrial network seems to be centered around the MTOC (Figure 7A; please see red arrows in Figure 7). In line with the live cell imaging data (Figure 6), the BCR clustered centrally and the mitochondrial network condensed around the MTOC-organized cluster upon BCR attachment (Figure 7C). Hence, we suggest that reorganization of the mitochondrial network correlates with elevated SPRC. The Seahorse analyses suggested – despite the lack of statistical significance – in every experimental setup that EFhd2KO B cells exhibit a higher SPRC than WT B cells on PDL (Figures 2-4). It was therefore intriguing to see that CD40/IL-4 activated EFhd2KO B cells attached to PDL already reveal a partial polarization of the mitochondrial network around loosely associated BCR clusters (Figure 7B), which did not connect properly to the MTOC, in line with the previous experiments (Figure 6B). This situation did not change obviously upon BCR attachment (Figure 7D). Rather unexpected, both BCR and mitochondrial networks emerged more dispersed and BCR clusters could not clearly be associated with the coating plane. These notions support the previous live cell imaging data (Figure 6). Next, we attempted to connect EFhd2 physically to the BCR synapse. Re-expression of EFhd2 in EFhd2KO B cells should i) restore mitochondrial organization and ii) disclose the localization of EFhd2 in primary B cells. Despite our experience in retroviral transduction of primary B cells, we were not able to express the EFhd2-EGFP fusion protein in primary B cells for unknown reasons (not shown). Therefore, we transduced EFhd2KO B cells with an HA-tagged EFhd2 in conjunction with IRES-GFP. Re-expression of EFhd2 conferred mitochondrial polarization and near substrate clustering in the BCR-attached transduced cells (Figure 7E). This experiment thereby confirms the direct function of EFhd2 in immune synapse organization. Figure S4 shows more examples of centralized mitochondria in the GFP^+^ EFhd2-transduced cells compared to the neighboring non-transduced (not green) ones. EFhd2 is present at the plasma membrane, in cell protrusions, cell-substrate contact sites and the EFhd2 network intercalates with the mitochondrial network. In sum, we identify here a mechanism which links antigen structure to mitochondrial fitness in activated B cells.

**Figure 7:**
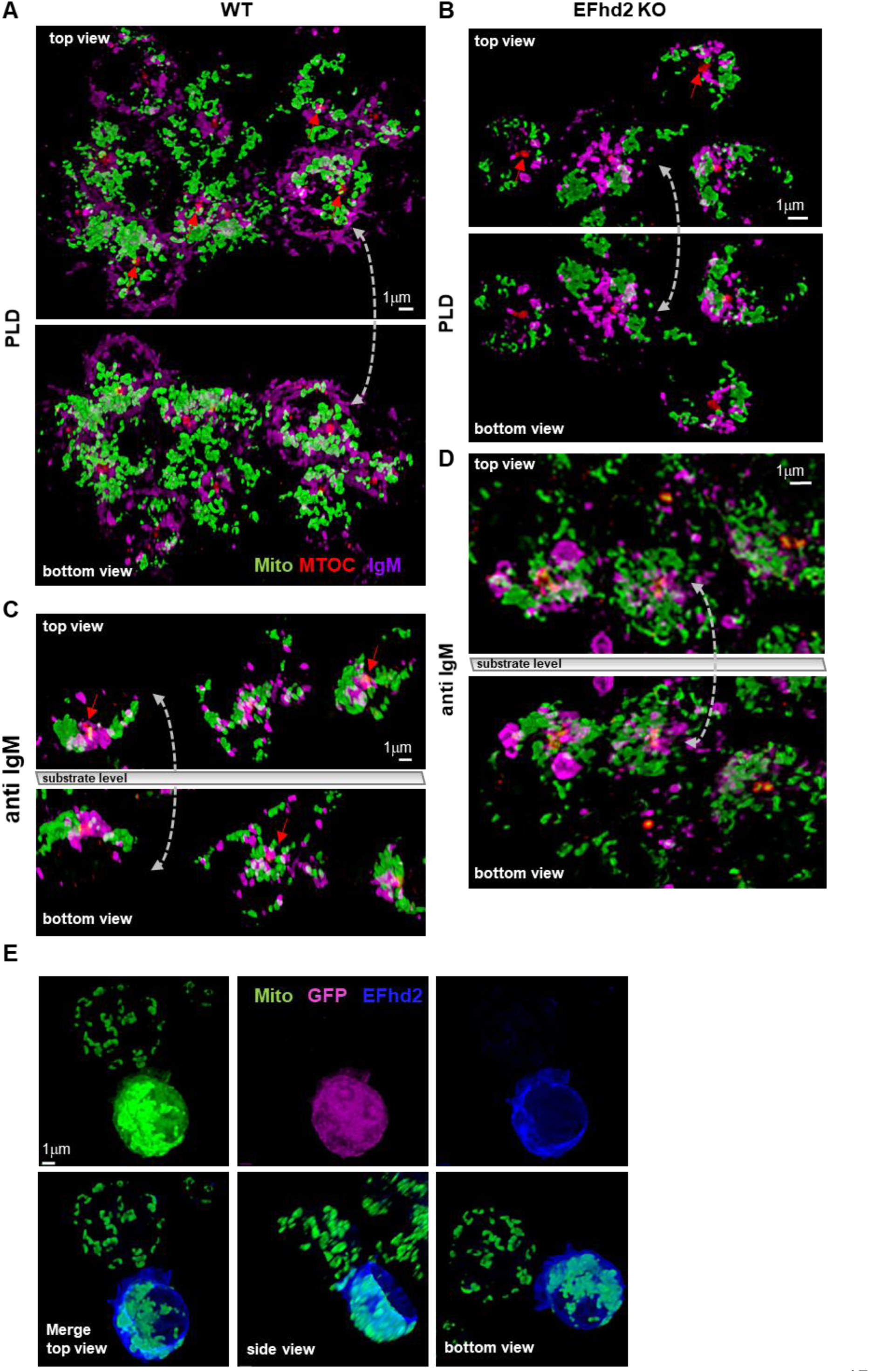
EFhd2 controls BCR and mitochondrial reorganization at the BCR synapse. CD40/IL-4 activated WT (A, C) and EFhd2KO B cells (B, D) were labelled with Mitotracker Green, attached to Poly-D-Lysine (PDL) or anti-IgM antibodies for ∼40 min, fixed, stained with anti-Tubulin (red) and anti IgM (purple) antibodies and analyzed by super resolution spinning disc confocal microscopy. Z-stacks were 3D reconstructed after deconvolution. Representative cells are shown from different angles. E. CD40/IL-4 activated EFhd2KO B cells were transduced to express EFhd2HA-IRES-GFP, attached to anti-IgM antibodies for ∼40 min, fixed and stained with anti HA-antibodies. Cells were analyzed by super resolution spinning disc confocal microscopy. Z-stacks were 3D reconstructed after deconvolution.EFhd2-expressing cells were identified by GFP fluorescence (false coloured, magenta, upper panel). Representative cells are shown from different angles without GFP fluorescence (lower panel). Scale bars, 1 μm.

## Discussion

This study demonstrates that B cells up-regulate mitochondrial metabolism upon BCR-synapse formation through EFhd2-mediated microtubule stabilization, which drives mitochondrial remodeling and supercharging at the synapse. This process may support catabolic stress resilience in activated B cells or LZ GC B cells awaiting co-stimulation before switching to anabolic pathways and clonal expansion ^18,44^. These findings could explain on the metabolic level why repetitive synthetic or natural antigen arrays, such as viruses ^4,45^, or immobilized antigen on (F)DC ^2,3,7^, elicit greater humoral immune responses. They may also be relevant for the activation of self-reactive B cells: polyvalent autoantigen complexes activate autoreactive B cells under physiological conditions and repeated encounter of autoantigen complexes leads to the production of affinity-matured autoreactive IgM ^46^. Because membrane-bound self-antigen can also elicit mitochondrial directed clonal deletion ^47,48^, balancing mitochondrial fitness and intrinsic apoptosis pathways could influence the fate of self-reactive B cells.

A major implication of this study is that mitochondrial function in B cells is not an isolated process governed by nutrient availability, mtDNA abundance ^26,28^ and pO_2_, but is tightly linked to cytoskeletal and membrane receptor dynamics. This integration ensures efficient ATP production and metabolic adaptability at the immune synapse, which is crucial for B cell survival and activation. Previous data indicate that the BCR elicits antigen polarization and mitochondrial reprogramming via Protein kinase C β ^49^. Here we follow up on these data to show that BCR polarization is directly linked to mitochondrial condensation at the BCR synapse. Further, we identify an essential player in this process, EFhd2. Our findings suggest that while EFhd2 is not essential for baseline mitochondrial function, it is crucial for dynamic metabolic adaptation during antigen recognition. This is especially important when cell-cell contacts are formed, like in the GC LZ, where EFhd2 abundance is highest. ^29^. These data suggest that EFhd2 serves as a molecular scaffold that integrates mitochondrial positioning with cytoskeletal organization. The mitochondrial reorganization correlated predominantly with SPRC under our conditions. A high mitochondrial SPRC equips cells with a high adaptive capacity towards stress and relies on a well-organized network of fused mitochondria. Vice versa, a low SPRC correlates with distorted mitochondria and sensitizes cells to metabolic stress ^50^. It can also be attributed to a mildly increased proton leak ^50^, which was reduced in BCR-attached but not in PDL-attached EFhd2KO B cells. Hence, we propose that EFhd2 controls mitochondrial positioning towards the BCR synapse via microtubule stabilization. This proposal aligns with the notion that mitochondrial transport is controlled by Kinesin activity and tubulin conformation ^51^. How EFhd2 controls microtubule turnover and mitochondrial dynamics in B cells remains to be resolved. However, in vitro data show that recombinant EFhd2 hinders Kinesin-mediated microtubule gliding^34^, pointing to a direct interaction of EFhd2 with either Kinesin or microtubules. In fact, in the hindbrain, EFhd2 interacts with proteins of the Actin, microfilament and tubulin cytoskeleton, as well as with proteins involved in mitochondrial regulation ^52^. In particular, there is an association between EFhd2, VPS13C and DNM1. VPS13C and DNM1 maintain mitochondrial membrane potential and mitochondrial fission in neurons, respectively ^52^. DNM1 is a key mediator of clathrin-dependent endocytosis. In line, in cancer cell lines EFhd2 functions as a cargo-specific adaptor in clathrin- and dynamin-independent endocytic pathways, facilitating the internalization and trafficking of various receptors, including adhesion receptors ^53^.

Several intracellular signaling pathways could mediate the effects of EFhd2 on mitochondrial and cytoskeletal organization. An obvious choice is the phosphatidylinositol-3-kinase-Akt-mTOR pathway, which guides metabolic reprogramming in B cells ^54^, yet we did not observe altered mTOR activity in BCR or anti CD40/IL-4 activated EFhd2KO B cells. As a sensor of the cellular energy status, AMP activated Kinase (AMPK) ^54^ might be aberrantly activated in EFhd2KO cells, altering cytoskeletal dynamics and mitochondrial bioenergetics, but in this case mTOR dysregulation would also be expected, which we did not observe. A clear candidate is altered Ca^2+^ signaling: EFhd2 has been implicated in the BCR-elicited cytoplasmic Ca^2+^ elevation in WEHI231 cells; unexpectedly, EFhd2 fulfils this function not in primary B cells ^30^ and unlike in WEHI231 cells, EFhd2 in primary B cells does not seem to control mitochondrial dysfunction. This difference to EFhd2-overexpressing WEHI231 cells could be a consequence of the fact that transformed B cells experience more mitochondrial stress due to their enforced cell cycle progression ^55^. However, EFhd2 does bind Ca^2+^ ^56,57^. It is possible that loss of EFhd2 modulates Ca^2+^ microislands in its vicinity and thereby affects Ca^2+^ based signaling pathways, such as PKCβ, which could impact mitochondrial function and cytoskeletal interactions ^49^. Whether and how the BCR regulates metabolic pathways via the cytoskeleton is unclear ^58^. Although we did not observe a function for EFhd2 in classical proximal signaling pathways ^30^ we do hypothesize that EFhd2 controls BCR signaling towards mitochondria via the cytoskeleton along the following arguments: i. an EFhd2-EGFP fusion protein localizes at the plasma membrane and in BCR clusters in transduced WEHI231 B cells ^35^; ii. destabilization of the Actin cytoskeleton by Latrunculin A activates BCR signaling via membrane remodeling ^59^; iii. EFhd2KO B cells show enhanced Actin remodeling ^29^; iv. activated EFhd2KO B cells appear to have already remodeled BCRs, mitochondria and enhanced SPRC, v. treatment of activated WT B cells with Latrunculin A heightens SPRC, vi. EFhd2 re-expression confers mitochondrial accumulation at the substrate. It remains to be determined whether EFhd2 connects the Actin and microtubule systems, or interacts with a protein fulfilling this function, perhaps in a Ca^2+^ dependent manner.

Owing to the complex interplay of signaling pathways and receptors on B cells we simplified the experimental setup, limiting ourselves to very rigid surfaces and in vitro cultures. However, positive B cell selection in the LZ involves FDC that provide a rather stiff membrane scaffold, enabling the specialized GC BCR synapse to exert force ^13^. Moreover, in vivo we did observe that EFhd2 supports B-FDC synapses in a B cell intrinsic manner and that EFhd2KO GC B cells vanish under competitive conditions ^29^. Hence, in light of these finding, we endorse the concept of EFhd2-mediated mitochondrial reorganization and fueling in the LZ that could support affinity maturation ^27^. This will be tested in future experiments. Notwithstanding, a limitation of this study is that it only addresses the IgM BCR and its capacity to discriminate between soluble and membrane bound stimulation via mitochondrial charging. It has been reported that the IgM BCR has a lower activation threshold than the IgD BCR and can respond to monovalent antigens ^60^. IgM and IgD receptors localize in different protein islands but IgD can interfere with IgM BCR signaling ^60,61^. Future experiments should also address the IgD BCR, which is no longer present in GC B cells, however, in GCs, IgG BCRs could also cluster mitochondria. Besides the BCR, the integrin leukocyte-function antigen 1 (LFA-1), binding to ICAM-1, and CXCR5, there are a number of other receptors on GC B cells that influence their fate, such as FAS ^62,63^. Non-apoptotic FAS signaling down-regulates mTOR activity in CD40 activated human B cells. This process involves regulation of EFhd2 ^63^. Moreover, EFhd2 does appear to modulate T cell activation differently upon soluble anti CD3/CD28 or bead-bound anti CD3/CD28 stimulation and is a component of the TCR synapse ^64,65^. Future experiments will show whether EFhd2-mediated mitochondrial organization has also relevance in other than B cells and identify other cytoskeletal key players and receptors in this process.

## Supporting information

Weckwerth_2025_Supplement

## Acknowledgments

This work was supported by grants from the Deutsche Forschungsgemeinschaft (DFG) GK2599 (to D.M.), FOR5560 (Mi939/7-1; to DM), Mi939/6-1, to D.M., the intramural Scholarship Programme ‘Promotion of Equal Opportunities for Women in Research and Teaching’ (FFL) of the Friedrich-Alexander-Universität Erlangen-Nürnberg (to L.W.) and the Core Unit Optical Imaging Center Erlangen (OICE). We thank the Core Unit Cell Sorting of the Friedrich-Alexander-Universität Erlangen-Nürnberg for excellent technical assistance.

## Materials & Methods

### Mice

The experiments were conducted using both female and male EFhd2-deficient mice (Brachs et al., 2014) on a C57BL/6 background. Wild-type littermates served as controls. All mice were housed under pathogen free conditions in the IVC at the Franz-Penzoldt-Center Erlangen (Germany). All experimental procedures were performed in accordance with ethical guidelines for animal experimentation and were approved by the government of Lower Franconia, Bavaria (Germany).

### Isolation of primary murine cells from spleen and mesenteric lymph nodes

Spleens and mesenteric lymph nodes were harvested and transferred into cold MACS buffer (Phosphate buffered saline (PBS), supplemented with 2% FCS). Tissues were gently passed through a 70 μm cell strainer (BD) using the plunger of a 5 mL syringe (BD). The resulting cell suspensions were pelleted by centrifugation at 400 x g for 5min at 4°C. For spleen samples, erythrocytes were lysed upon resuspension in red blood cell-lysis buffer (150mM NH_4_Cl, 10mM KHCO_3_ and 1mM EDTA) for 5min at room temperature. The reaction was stopped by adding cold MACS buffer, followed by centrifugation at 400 x g for 5min at 4°C. The final cell suspensions were filtered through a 30µm mesh filter (Sysmex) and maintained in cold MACS buffer until further use.

### Purification and in vitro cultivation of primary murine B cells

Splenic B cells were enriched using the EasySep Mouse B cell isolation negative selection kit (StemCell Technologies, EasySep #19854) according to the manufacturer’s instructions. Briefly, spleen cells were resuspended in MACS buffer, surface blocked with rat serum, and immunomagnetically enriched for untouched naive B cells. The purity of the isolated B cells was assessed by surface staining for CD19 and B220. Usually, an enrichment of > 95% was achieved. Freshly isolated B cells were immediately cultured at a concentration of 2×10^6^ cells/ml in pre-warmed R10 medium (RPMI1640 supplemented with 10% fetal calf serum [FCS], 2mM glutamate, 1mM sodium pyruvate, 50U/mL penicillin G, 50μg/mL streptomycin and 50μM ß-Mercaptoethanol) for 24-48h at 37°C and 5% CO_2_. Cultures were supplemented with 10µg/ml anti-IgM F(ab’)_2_ Ab fragment (Jackson ImmunoResearch), 10µg/ml anti-CD40 Ab (clone FGK45, Protein G purified from hybridoma supernatants) and 0.1U/ml IL-4 (mouse IL-4, Miltenyi Biotec).

### Retroviral infection

The plasmid pCru_EFhd2Myc_IRES-GFP ^35^ was used to generate pCru_EFhd2HA_IRES-GFP. PlatinumE packaging cells were transfected with plasmid, retroviral supernatant was collected after 48 and 72h, 0.45 μM filtered and stored at −70°C. 10^6^ 24h activated B cells were incubated with 1 ml of retroviral supernatant containing 4μg Polybrene, centrifuged for 3h at 3300rpm, 33°C in a Heraeus Tabletop centrifuge, washed once in fresh medium and further grown in the appropriate stimulant.

### Measurement of mitochondrial mass and membrane potential

Mitochondrial mass and membrane potential were assessed by staining cells with Mitotracker Green and DeepRed (Thermo Scientific), respectively, followed by flow cytometric analysis. For staining, 1×10^6^ – 4×10^6^ cells were resuspended in pre-warmed freshly made Mitotracker staining buffer (RPMI without supplements, 10nM Mitotracker Green and 5nM Mitotracker DeepRed; 100µl per 1×10^6^ cells). After incubation at 37°C and 5% CO_2_ for 30min, cells were washed twice in RPMI and blocked with 50µl unlabeled anti-CD16/32 Ab (10 µg/mL in RPMI) for 15 min on ice. Cells were then washed once by centrifugation at 400 x g for 5min at 4°C, resuspended in 50µl RPMI containing the appropriate fluorochrome-conjugated Abs and incubated for 20min on ice in the dark. Afterwards, cells were washed twice with PBS and immediately analyzed using a Gallios flow cytometer (Beckman Coulter). Data analysis was performed using Kaluza software version 2.1. For intracellular cell staining, cells were permeabilized and fixed using the FoxP3/Transcription Factor Staining buffer set (eBioscience) according to the manufacturer’s instructions.

### Extracellular flux assay

Extracellular flux analysis was performed using a standardized protocol to measure the oxygen consumption rate (OCR) on a Seahorse XF 96 Analyzer (Agilent). Briefly, cartridges were calibrated overnight with 200µl/well calibrant buffer (Agilent). Cell culture plates were coated with 100µl/well Poly-D-Lysine (100µg/ml in 1x Tris EDTA buffer) at room temperature (RT) for 1-2h, or with anti-IgM F(ab’)_2_ Ab fragment (as indicated, usually 10 μg/ml, in 1x TE buffer) over night at 4°. Plates were then dried for 1h at RT, and washed three times with ddH_2_O to remove unbound antibodies. Cells were resuspended in the respective Seahorse XF RPMI assay medium (supplemented with 1mM pyruvate, 2mM glutamine and 10mM glucose for a Mito Stress Test) and seeded at a density of 2.5 x 10^5^ per well in 180µl. Measurements were performed at least in triplicates. To ensure proper attachment and even distribution, cells were briefly centrifuged. For plating on anti-IgM-coated plates, an additional incubation step for 0.5– 4 hours at 37°C and 5% CO₂ was performed. Cells were treated sequentially with Latrunculin A (1µM), Nocodazole (0.1µM) or Paclitaxel (1µM) or left untreated, and then starved for 45min at 37°C in a non-CO_2_ incubator. The Mito Stress Test assays were performed according to the manufacturer’s instructions. Viability before and after the experiments was routinely assessed by propidium iodide exclusion in an automated cell counter (Nucleocounter NC-300, Chemometec).

### Spinning Disc live cell confocal microscopy

Live cell microscopy was performed at 37°C and 5% CO_2_ using the Evident Super Resolution Spinning Disc microscope (LSM) with a 63x oil immersion objective. 8-well polymer chambered coverslips (Ibidi) were coated with either anti-IgM F(ab’)_2_ Ab fragment (10µg/ml in PBS) or poly-D-Lysine (100µg/ml in PBS) overnight at 4°C. Chambers were then blocked with 200μl/well of 10mg/ml fatty acid free BSA in PBS for 2h at room temperature, followed by washing with pre-warmed PBS. Activated B cells were stained with different loading dyes (1µM Mitotracker Green and 1µM Tubulintracker DeepRed in RPMI) for 30min at 37°C and 5% CO_2_ and with anti-IgM F(ab’)_2_ Ab (AF 594, diluted 1:100 in RPMI, Jackson ImmunoResearch) for 20min on ice in the dark. After staining, cells were washed twice with RPMI and seeded at a concentration of 5×10^5^ cells/ml in 200µl/well. Fluorescence signals were detected immediately after seeding at appropriate wavelengths using standard filters (e.g., FITC, Cy5, DAPI and Rhod). Signals and differential interference contrast (DIC) were recorded with an Evolve camera. Image acquisition was performed using the Zeiss Zen software. Fiji was used to further analyse the images; the MTOC-center distances and the weighted MTOC-BCR distances were quantified using custom Fiji macros. Further image processing was carried out using Fiji software and the 3Dscript plugin ^66^.

### Spinning Disc super resolution confocal microscopy

8-well polymer chambered coverslips (Roth or Thermo Scientific) were coated with either anti-IgM F(ab’)_2_ Ab fragment (10µg/ml in PBS) or poly-D-Lysine (100µg/ml in PBS) overnight at 4°C. Activated B cells were stained with 1µM Mitotracker Green for 30min at 37°C and 5% CO_2_ for 20min on ice in the dark. After staining, cells were washed twice with RPMI and seeded at a concentration of 1×10^6^ cells/ml on the wells. After 40 min, cells were fixed in 4% paraformaldehyde at 37°C for 15 min, washed in PBS, permeabilized for 5 min with 0.1% Triton X-100 and blocked with 10% FCS in PBS for 30 min. Cells were incubated with goat anti-IgM antibodies coupled to Alexa647, washed in PBS and fixed in 1% PFA for 15 min. Afterwards, cells were stained with rabbit anti Tubulin antibodies, followed by goat anti rabbit antibodies conjugated to Alexa Fluor 594. In some experiments, Mitotracker Green labeled cells were fixed, permeabilized and stained with rat anti HA antibody coupled to Alexa 647. After washing in PBS, slides were mounted with Mowiol. Fluorescence signals were detected at appropriate wavelengths using standard filters. Z-stacks (310 nm per slice) were recorded on an Olympus IXplore Spin SR microscope with a 100x glycerin objective. Z-stacks containing 10-20 layers were deconvolved using Evident cellSense software. Z-stacks were 3D reconstructed using Fiji software and the 3Dscript plugin ^66^.

### Cell apoptosis assay – IncuCyte

Activated B cells were plated at a concentration of 4×10^5^ cells/ml in 50µl RPMI medium (supplemented with the respective stimulants) on 96 well plates coated with either Poly-D-Lysine (100µg/ml in PBS) or anti-IgM F(ab’)_2_ Ab (10µg/ml in PBS). Plates were washed once to remove unbound Ab. Cells were imaged every 1 to 2 hours using the IncuCyte Zoom Live-Cell Analysis System (Essen Bioscience). For cell death assays, 250nM IncuCyte Cytotox green fluorescent dye (Sartorius) was added to the cells. Apoptosis was calculated by counting green objects and normalizing the count to the confluence factor and the initial time point.

### Reagents and antibodies

| REAGENT or RESOURCE | SOURCE | IDENTIFIER |
| --- | --- | --- |
| **Antibodies Mouse** |  |  |
| Anti B220 (APCFire750, clone Ra3-6b2) | Biolegend | Cat# 103278 |
| Anti CD16/32,clone93 | Biolegend | Cat# 101302 |
| Anti CD19 (APCFire750, clone 6D5) | Biolegend | Cat# 115558 |
| Anti CD19 (BV421, clone 6D5) | Biolegend | Cat# 103278 |
| Anti CD38 (PerCPCy5.5, clone 90) | Biolegend | Cat# 102722 |
| Anti CD95 (PE.Cy7, clone Jo2) | Biolegend | Cat# 557653 |
| Anti GL-7 (Pacific Blue, clone GL-7) | Biolegend | Cat# 144613 |
| Anti IgM F(ab')<sub>2</sub> fragment (AF594) | Jackson<br>ImmunoResearch | Cat# 115-587-020 |
| Anti pS6 (PE, clone cupk43k) | eBioscience | Cat# 12-9007-42 |
| **Tracker** |  |  |
| MitoTracker DeepRed | Thermo Fisher<br>Scientific | Cat# M22426 |
| MitoTracker Green | Thermo Fisher<br>Scientific | Cat# M7514 |
| Incucyte® Cytotox Green Dye | Sartorius | Cat# 4633 |
| TubulinTracker DeepRed | Thermo Fisher<br>Scientific | Cat# T34077 |
| **Experimental Models: Organisms/Strains** |  |  |
| EFhd2 KO mice | Brachs et al., 2014 | MGI#5707812,RRID:MGI:5707812 |
| **Chemicals, Peptides, and Recombinant Proteins** |  |  |
| Anti CD40,cloneFGK45, | Rolinketal.,1996 | RRID:AB_2490239 |
| Antimycin A | Sigma | Cat# A-8674 |
| β-Mercaptoethanol | Gibco - Thermo<br>Fisher Scientific | Cat# 31350-010 |
| Carbonyl cyanide-p-trifluoromethoxy-phenylhydrazone (FCCP) | Sigma | Cat# C2920 |
| CellCount and Viability Solution 13 | Chemometec | Cat# 910-3013 |
| Fetal bovine serum (FCS), heat-inactivated | Gibco | Cat# 10270-10 |
| Glucose | Carl Roth | Cat# HN06.3 |
| Goat anti-IgM AffiniPure F(ab')<sub>2</sub> $\mu$ chain specific | Jackson<br>ImmunoResearch | Cat# 115-006-075 |
| Interleukin-4, premium grade | Miltenyi Biotec | Cat#130-097-761 |
| Latrunculin A | Sigma | Cat# 428026 |
| L-glutamine | Gibco - Thermo<br>Fisher Scientific | Cat# 25030-24 |
| Nocodazole | Sigma | Cat# 487929 |
| Oligomycin | Sigma | Cat# 75351 |
| Paclitaxel | Sigma | Cat# T7191 |
| Penicillin/Streptomycin | Gibco - Thermo<br>Fisher Scientific | Cat# 15140-122 |
| Poly-D-Lysine | Sigma | Cat# P7280 |
| Rotenone | Sigma | Cat# R8875 |
| RPMI1640 | Gibco - Thermo<br>Fisher Scientific | Cat# 31870-25 |
| Sodium Pyruvate | Gibco - Thermo<br>Fisher Scientific | Cat# 11360-039 |
| XF Calibrant buffer | Agilent | Cat# 100840-000 |
| XF RPMI Assay medium [pH 7,4] | Agilent | Cat# 103576-100 |
| 4-Hydroxy-3-nitrophenylacetyl (bovine serum albumin; BSA), fatty acid free | Sigma | Cat# A3803 |
| **Software and Algorithms** |  |  |
| GraphPad Prism 9 | GraphPad | https://www.graphpad.com/scientific-software/prism/ |
| Kaluza | Beckman Coulter | https://www.beckman.de/flow-cytometry/software/kaluza |
| Seahorse Wave Desktop Software | Agilent | https://www.agilent.com/en/product/cell-analysis/real-time-cell-metabolic-analysis/xf-software/seahorse-wave-desktop-software-740897 |
| **Others** |  |  |
| EasySep™ Mouse B Cell Isolation Kit | StemCell | Cat# 19854 |
| NucleoCounter NC-3000 | ChemoMetec | N/A |
| μ slide 8-well polymer chambered coverslips | Ibidi Bioscience | Cat# 80821 |

## Abbreviations

Ab: antibody
APC: antigen presenting cell
ATP: adenosine triphosphate
BCR: B cell receptor, electron transport chain
FCCP: carbonyl cyanide m-chlorophenyl hydrazine
FDC: follicular dendritic cell
FL: fluorescence
FSc: Forward Scatter
GC: germinal center
GFP: green fluorescent protein
ICAM-1: intercellular adhesion molecule
IL-4: Interleukin-4
KO: knock-out
LatA: Latrunculin A
LFA-1: leukocyte-function antigen
MBC: memory B cell
MT: microtubule
MST: mito stress test
MTOC: microtubule organizing center
Noc: Nocodazole
PDL: Poly-D-Lysin
PTX: Paclitaxel
OxPhos: oxidative phosphorylation
OCR: oxygen consumption rate
Ψ: mitochondrial membrane potential
SHM: somatic hypermutation
SSc: Side scatter
WT: wildtype

