## Supplementary material for "Swiprosin-1/EFhd2 promotes mitochondrial spare capacity in response to immobilized antigen in B cells via microtubule stabilization": Weckwerth_2025_Supplement

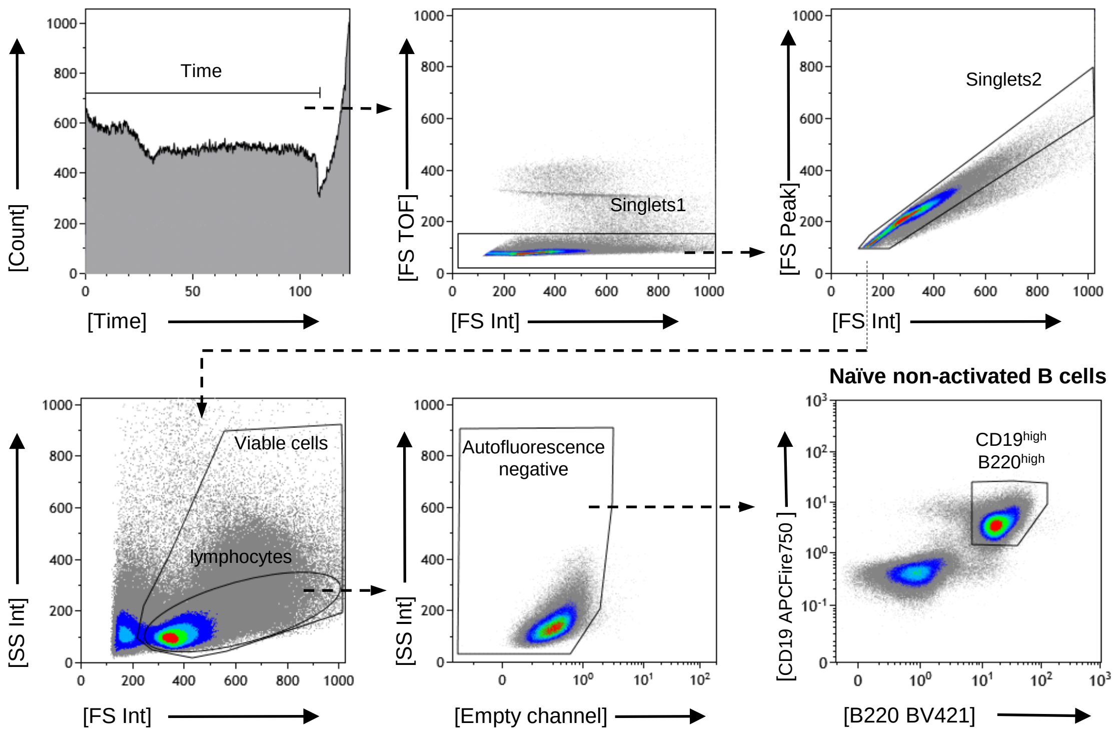


**Figure S1**
**Flow cytometric gating strategy**

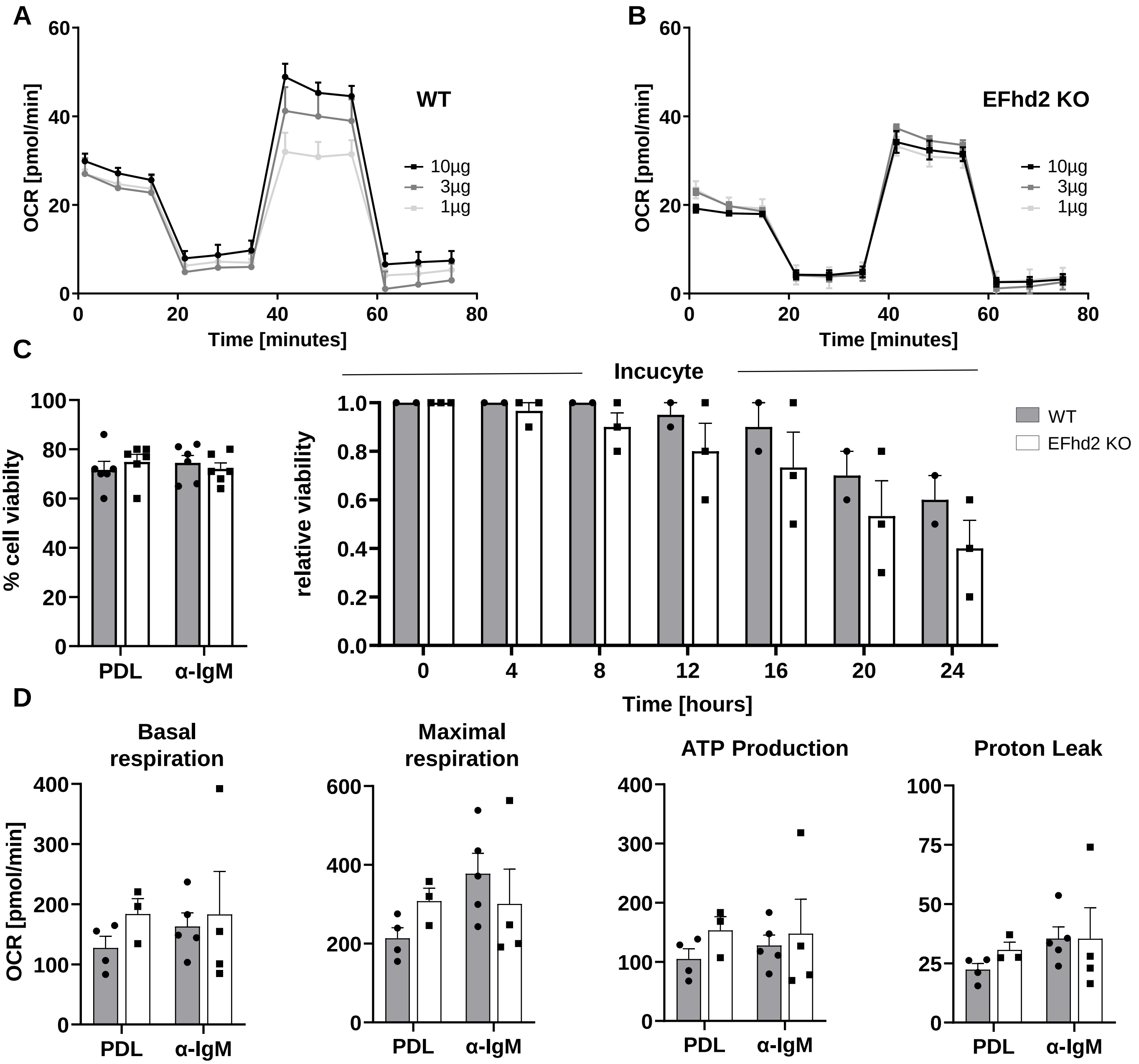


**Figure S2.** A, B. Representative extracellular flux analysis of 24h anti-IgM Ab activated splenic B cells attached to anti IgM antibodies with different coating concentrations. The basal oxygen consumption rate (OCR) was measured before and after injection of oligomycin, FCCP and rotenone plus antimycin A using the Seahorse Wave Mito Stress Test (MST) protocol. Symbols represent means of two mice, each of three to four replicate wells, mean ± SEM. C. Viability of of 24h anti-IgM Ab activated splenic B cells attached to PDL or anti IgM antibodies after the Seahorse Mito stress test or in an Incucyte chamber. D. Calculated OCR of basal respiration, maximal respiration, ATP production and proton leak of 24h anti-CD40/IL-4 activated B cells attached to PDL or anti IgM antibodies.

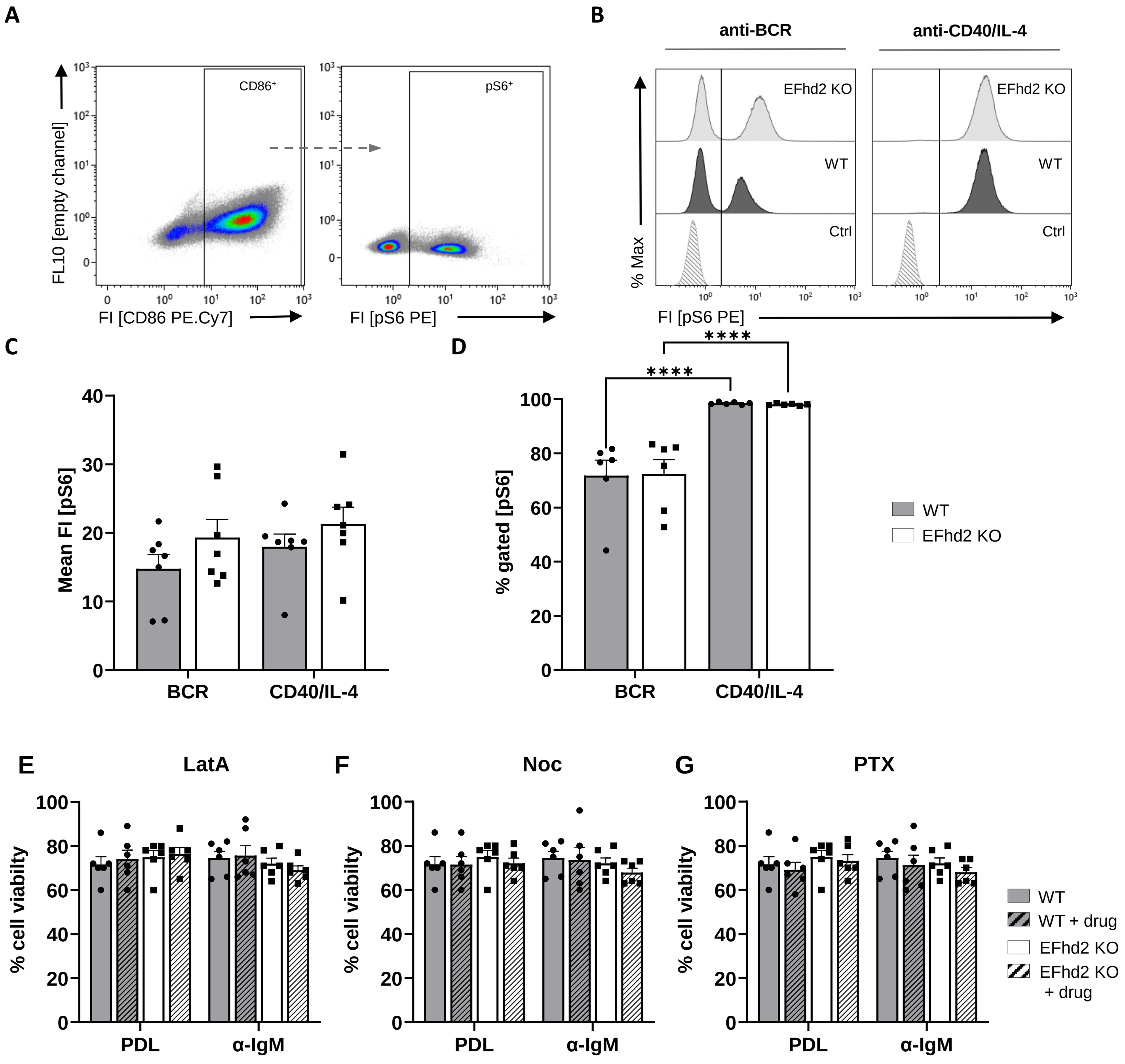


**Figure S3 Mtor activity and viability in activated WT and EFhd2KO B cells**

A. Splenic B cells from wild type (WT) and EFhd2 KO mice were stimulated with anti-IgM (BCR) or anti-CD40/IL-4 for 24h. Membrane staining for CD86 was performed as a marker of activation. B. The cells were then fixed, permeabilized and stained with anti-pS6 antibodies, ctrl.: unstained. C. Statistical analysis of the MFIs and D. frequencies of pS6$^{+}$ WT and EFhd2 KO B cells. Data are presented as mean ± SEM; one data point ≙ one mouse; statistics: N=2, n=6 significance was calculated using Two-way ANOVA (p values <0.5. *p < 0.05, **p < 0.01, ***p < 0.002, ****p < 0.0004). E-F: Viability of of 24h anti-IgM Ab activated splenic B cells attached to PDL or anti IgM antibodies after the Seahorse Mito stress test. c - f: data are presented as mean ± SEM; each dot represents one mouse; statistics: N=1-2, n=3-6; significance was analyzed using Two-way ANOVA (p values <0.5. *p < 0.05, **p < 0.01, ***p < 0.002, ****p < 0.0004).

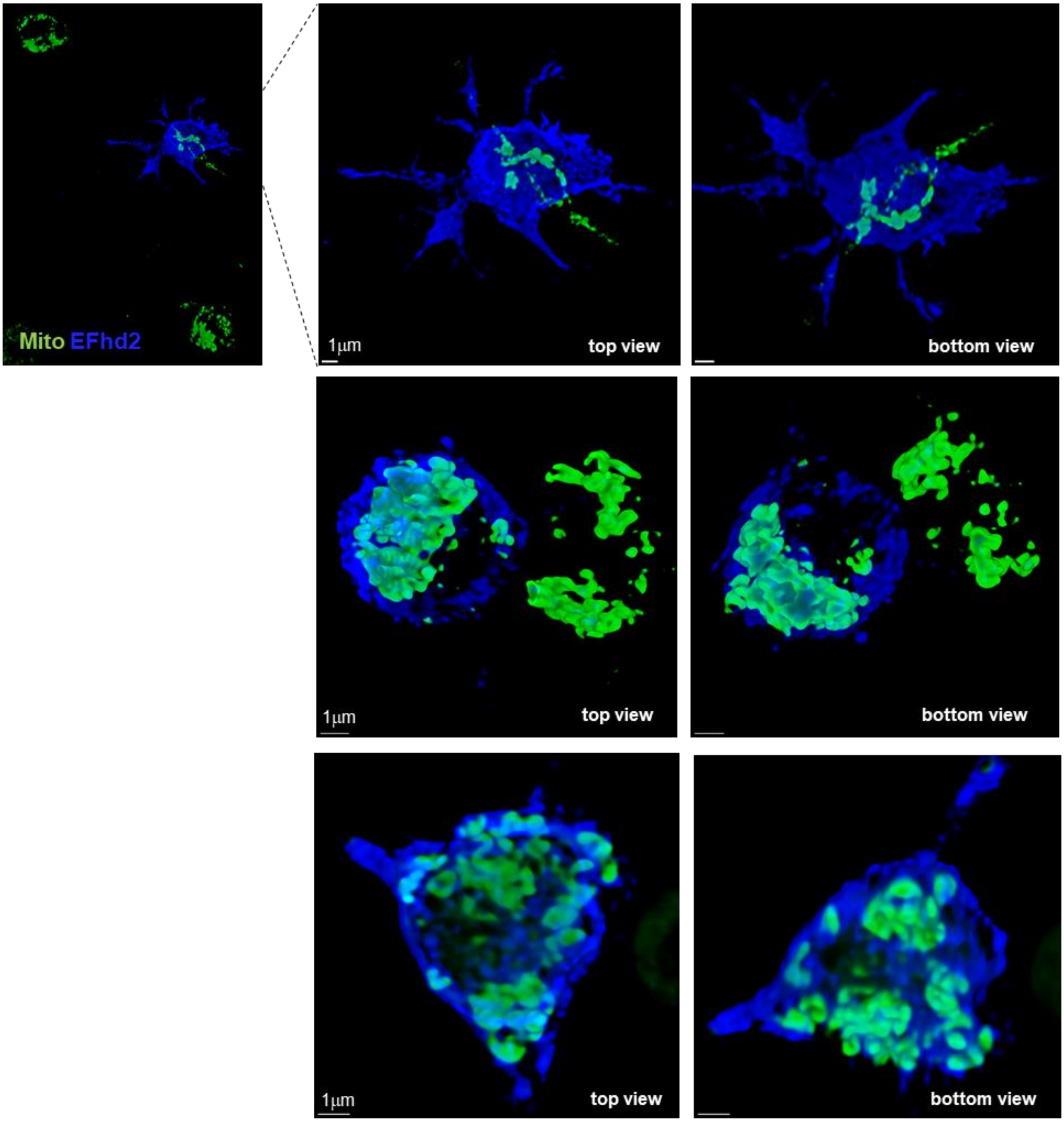


**Figure S4 Localization of EFhd2 and mitochondria in EFhd2-transduced EFhd2KO B cells**
CD40/IL-4 activated EFhd2KO B cells were transduced to express EFhd2HA-IRES-GFP, attached to anti-IgM antibodies for ~40 min, fixed and stained with anti HA-antibodies. Cells were analyzed by super resolution spinning disc confocal microscopy. Z-stacks were 3D reconstructed after deconvolution. Representative cells are shown from different angles. EFhd2-expression was identified by GFP fluorescence (not shown here, but compare Figure 7). Scale bars, 1 μm.
